# Redox state changes of mitochondrial cytochromes in brain and breast cancers by Raman spectroscopy and imaging

**DOI:** 10.1101/2020.05.08.083915

**Authors:** Halina Abramczyk, Jakub Maciej Surmacki, Beata Brozek-Pluska

## Abstract

This paper presents a non-invasive approach to study redox status of cytochromes *in vitro* human brain cells of normal astrocytes (NHA), astrocytoma (CRL-1718), glioblastoma (U87- MG) and medulloblastoma (Daoy), and human breast cells of normal cells (MCF10A), slightly malignant cells (MCF7) and highly aggressive cells (MDA-MB-231), in vivo animal models, and *ex vivo* brain and breast tissues surgically resected human specimens by means of Raman microspectroscopy at 355 nm, 532 nm, 785 nm and endospectroscopic Raman probe at 785 nm. Here we show that the amount of reduced cytochrome becomes abnormally high in human brain tumors and breast cancers. In contrast, the amount of reduced cytochrome c is lower in cancer cells when compared to the normal one at *in vitro* conditions when the effect of microenvironment is eliminated. Mitochondrial dysfunction and alterations in the chemical composition of the nucleus, mitochondria, lipid droplets, cytoplasm in single cells have been detected by Raman imaging. Incubation *in vitro* with retinoic acid increases the amount of reduced cytochrome c.

## Introduction

In recent years, growing interest toward tumor redox status related to cytochrome family has highlighted as an important factor for cancer progression, resistance to treatments, and a poor prognosis. Cytochrome family (cytochromes b, c, a) has been shown to be up-regulated in many types of cancers.^1^ However, its role in brain and breast cancers has not been well understood.^2^ Cytochrome c (Cyt c) is an electron carrier protein that localizes in mitochondrion intermembrane space that transfers electrons from heme c1 in complex III to heme a3 in complex IV. Mitochondrial Cyt c has been identified to have dual function in controlling both cellular energetic metabolism and apoptosis and is one of the most intensively studied proteins.

Separately conventional cancer biology with genomic, transcriptomic and proteomic protocols can provide only a partial picture of cancer redox status. To be able to capture an idea of redox balance in normal cells and its aberrations in cancer development, progression, and proliferation we need new tools to monitor biochemistry, properties of single normal and cancer cells, tissues in animal models and in humans. Raman spectroscopy offers the ability to probe biochemical changes in mitochondria, lipid droplets, nuclei, cytoplasm, membrane and visualize the complex molecular events directly in living normal and cancer cells in vitro, ex vivo human tissues and in vivo animal models.^3,4^ It is also demonstrated as a versatile clinical diagnostic tool with numerous successful reports on the detection of cancerous tissues in human patients.^5–8^

Our goal was to demonstrate possibility of monitoring of redox state changes occurring in mitochondrial b- and c-type cytochromes in cancers. A thorough understanding of cytochrome role in brain and breast cancers with our new method will help establish Raman spectroscopy as a competitive clinical diagnosis tool for cancer diseases involving mitochondrial dysfunction.

## Results

To properly address redox state changes of mitochondrial cytochromes in brain and breast cancers by Raman spectroscopy and imaging, we systematically investigated how the Raman method respond to in vitro cells, animal models, ex vivo human tissues.

Fig.1A shows the first in Europe Raman- guided in vivo brain analysis of the brain for the animal model – living rat – using Raman endospectroscopy performed by Abramczyk et al.^4^ The first Raman- guided in vivo human brain analysis was performed by Desroches et al. in 2019.^9^ Fig.1B presents the Raman spectra of living rat brain obtained by using the hand-held Raman spectrometer with endospectroscopic fiber probe in the Raman guided in vivo measurement for normal brain at 785 nm and compared with the ex vivo normal brain tissue surgically resected specimens of animal model (rat) at the excitation 532 nm and at 785 nm.

**Fig.1.**
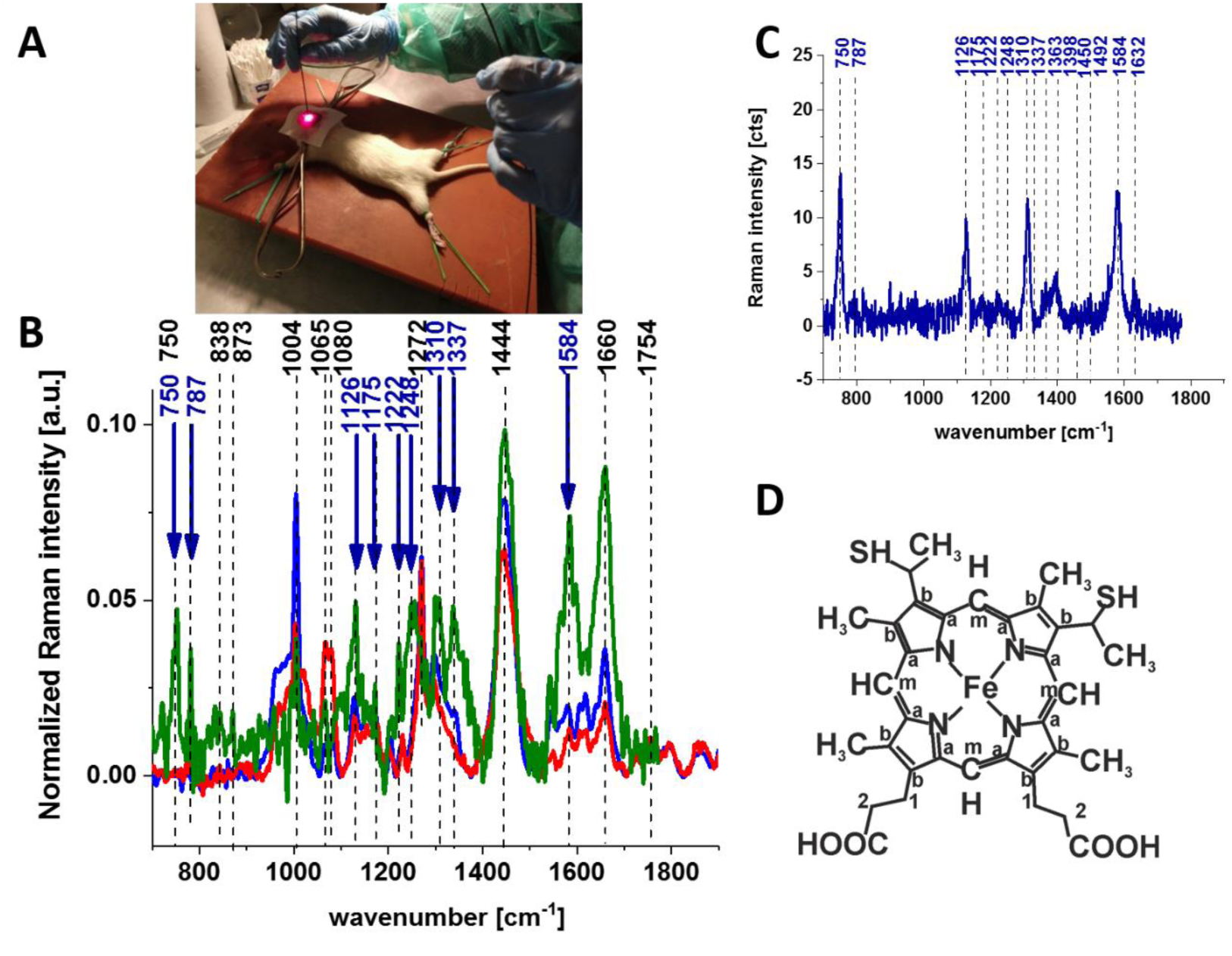
Raman- guided in vivo animal (rat) brain analysis (A), the average (n=6) Raman spectrum of the in vivo brain of animal model (rat) at the excitation 785 nm ▬ and of the ex-vivo brain of animal model (rat) at the excitation 532 nm ▬ and 785 nm ▬ (B), Raman spectrum of cytochrome c ▬ (dark blue) (C), structural formula of heme c in cytochrome c (D).

The relative intensities of Raman bands in Fig.1B depend on the excitation laser wavelength which clearly indicates that Resonance Raman effect is responsible for these differences.

Since 532 nm laser excitation causes Raman scattering from c and b-type cytochromes due to electronic absorption of cytochromes family (Q bands at 500-550 nm related to intraporphyrin transitions of the haem group in Cyt c)^10,11^ cytochromal Raman peaks may originate from both cytochrome types. Comparison of the Raman spectra of the brain tissue and purified cytochrome c in Fig.1C shows that the Raman enhancement upon 532 nm excitation corresponds perfectly to vibrations of Cyt c.^12,13^ However, literature assignments^14–17^ show that some cytochromal peaks are common to c, c1 and b cytochromes corresponding to vibrations of Cyts c, c1 or b. Thus, the major peaks at 750 and 1126 cm^-1^ are present in both types of cytochromes, whereas the peak at 1310 cm^-1^ corresponds to c-type cytochromes and the peaks at 1300 and 1337 cm^-1^ — to b-type cytochromes. The peak at 1337 cm^-1^ can be useful to distinguish cytochrome c and b, as the vibration at 1337 cm^-1^ represents a unique peak of the reduced Cyts.b (ferro- (Fe^2+^) cytochrome). Therefore, peaks at 750 and 1126 cm^-1^ in Raman spectra of the brain tissue in Fig.1 represent c, c1 and b-types of cytochromes. However, relative contributions of these peaks to the overall Raman spectrum differ in biological systems.^14–17^

The vibrations of cytochrome c that are enhanced in the brain tissue are presented in Fig.1B (blue color arrows). We observe five intensive peaks: 750 (symmetric vibrations of pyrrole rings), 1126 (vibrations of C_b_-CH_3_ side radicals), 1310 (vibrations of all heme bonds), 1363 (mode (ν_4_)) and 1584 cm^-1^ (v_19_ mode, vibrations of methine bridges (C_α_C_μ_, C_α_C_m_H bonds) and the C_α_C_β_ bond). There are also a number of other peaks with lower intensities 1248, 1352, 1632 cm^-1^ (methine bridges (bonds C_α_C_m_, C_α_C_m_H)). The Raman bands of the reduced form have higher intensities.^13^ Symmetric vibrational modes of the porphyrin ligand in Cyt c are enhanced to a greater degree using excitation wavelengths within the Soret absorption peaks at 408 nm (ferric Fe^+3^), 416 nm (ferrous Fe^+2^) states (see Fig.S1 in Supplementary Material), whereas asymmetric modes are enhanced to a greater degree using excitation wavelengths within the Q absorption peak at 500-550 nm.^18^ Detailed vibrational assignment can be found in ref^13^ Q-resonant Raman spectra contain unusually strong depolarized bands. In fact, the B_1g_ pyrrole breathing mode v_15_ (750 cm^-1^) gives rise to one of the strongest bands (Fig.1C). The bands of cytochrome c at 750, 1126, 1248, 1310, 1363 cm^-1^ are depolarized and represent the reduced form. Anomalously polarized bands appear in the Q-resonant spectra. Especially striking is the v_19_ mode^13^ (1584 cm^-1^), which produces one of the most prominent bands in the perpendicularly polarized psectrum. The band at 1584 cm^-1^ represents the reduced form of cytochrome and it is not observed in the oxidized form. Some of the peaks of the oxidized form of Cyt c (around 750, 1130, 1172, 1314, 1374, 1570-1573 and 1639 cm^-1^) have the same positions as the reduced form^19^, but their intensities are significantly lower.^11,19–21^

In the view of the results for animal brain it would be extremely valuable to control cytochrome activity in humans. To help address these challenges Fig.2A shows Raman enhancement of cytochromes for ex vivo human brain tissue of highly aggressive medulloblastoma, where the blue arrows shows the Raman peaks of Cyt c (reduced form). Note that exactly the same as in Fig.1 Raman enhancement is observed for the human brain at 532 nm. The most significant resonance Raman enhancement at 532 nm is observed for the reduced forms of Cyt c at 1584 cm^-1^. Details of this enhancement will be discussed later in this paper. However, there are also striking differences between the Raman spectra at 532 nm and 785 nm excitations compared to the results in Fig.1. Fig.2A reveals resonance Raman enhancement of some vibrations at 785 nm when the Cyt c resonance enhancement disappears. Some of the enhanced vibrational modes corresponds to tyrosine and phosphorylated tyrosine.^22^ The effect was weaker for the animal brain in Fig.1. The main reason for these differences may be related to the fact that Fig.1 shows normal brain tissue whereas Fig.2 shows the tumor brain tissue (medulloblastoma). This point will be discussed later when we will attempt to rationalize the whole picture that is revealed from our measurements.

**Fig.2.**
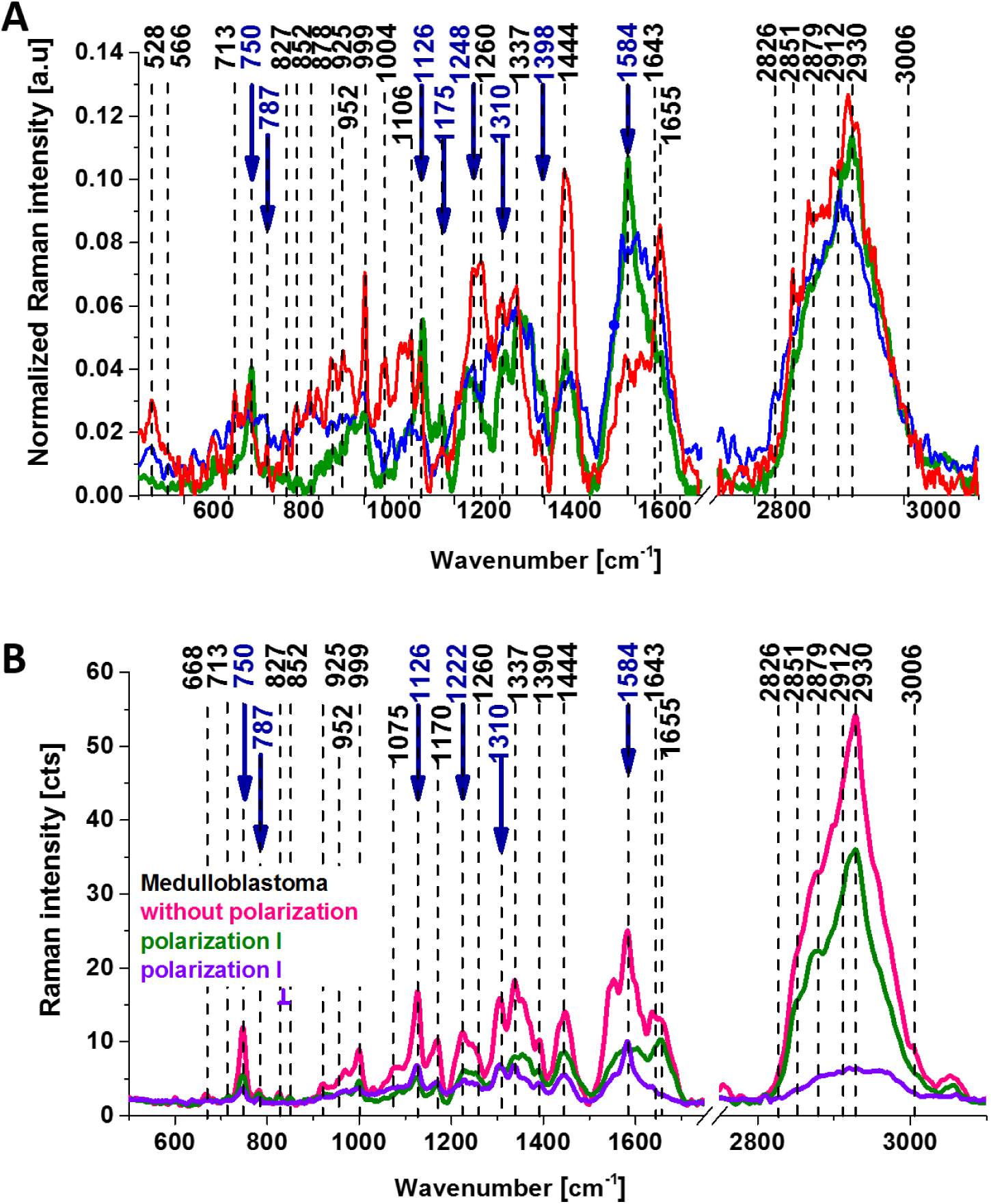
The Raman spectra for the human brain tissue of medulloblastoma at different excitations and experimental geometries for Raman scattering. (A) The average (n=6) Raman spectrum of the ex vivo tumor human brain tissue of medulloblastoma surgically resected specimens at the excitations 355nm ▬ (blue), 532 nm ▬ (green) and 785 nm ▬ (red). (B) The Raman spectra for the human brain tissue of medulloblastoma at 532 nm at different experimental geometries for Raman scattering: without polarization analyzer ▬ (magenta), at parallel ▬ (green) and perpendicular ▬ (violet) polarizations of the incident and Raman scattered beams.

As the cytochrome and tyrosine activity in normal and cancer cells may be related to the protein induced distortions and conformational changes we employed the polarized Raman spectroscopy. Polarized Raman spectroscopy^23^ provides vital information about a molecular structure, orientation in highly ordered systems and conformational preferences of a specific crystalline structure. Fig.2B shows the Raman spectra for the human brain tissue of medulloblastoma at different experimental geometries for Raman scattering: without a polarization analyzer, and at parallel I_II_ and perpendicular polarizations I_┴_ of the incident and Raman scattered beams. One can see from Fig.2B that the vibrational mode at 1584 cm^-1^ shows anomalously polarized band appearing in the Q-resonant spectra at 532 nm which produces one of the most prominent bands in the perpendicularly polarized spectrum. The anomalously polarized band at 1584 cm^-1^ appearing in the Q-resonant Raman spectra is characteristic of Cyt c.^13^ This finding additionally supports our previous conclusion that the Q-resonant Raman band at 1584 cm^-1^ in the brain tissue represents Cyt c.

Since Raman scattering at 1584 cm^-1^ is not observed in the oxidized form of Cyt c at 1584 cm^-1^ originates from the reduced form of Cyt c.^21^ Our results therefore suggest that the Raman intensity of 1584 cm^-1^ peak is the most sensitive vibration of redox status in the cell and is related to the amount of reduced Cyt c. Therefore, we used it to study the redox state of mitochondrial Cyt c in brain cells in vitro, ex vivo human tissues, in vivo animal models (rat).

To check whether the redox state of Cyt c is related to the cancer aggressiveness we used the Raman redox state biomarker represented by the Raman intensity of 1584 cm^-1^ peak.

Based on the average values (n= 44) obtained for the Raman biomarker of Cyt c I(1584/1444), (the ratio of the Raman intensities at 1584 and 1444 cm^-1^) we obtained a plot as a function of tumor grade malignancy. Fig.3 shows the Raman biomarker I(1586/1444) as a function of breast cancer grade malignancy G0-G3 (Fig.3A) and brain tumor grade malignancy G0-G4 (Fig.3B) for the human tissues and for in vitro human brain cells of normal astrocytes (NHA), astrocytoma (CRL-1718), glioblastoma (U87-MG) and medulloblastoma (Daoy), and human breast cells of normal cells (MCF10A), slightly malignant cells (MCF7) and highly aggressive cells (MDA-MB-231) (B).

**Fig.3.**
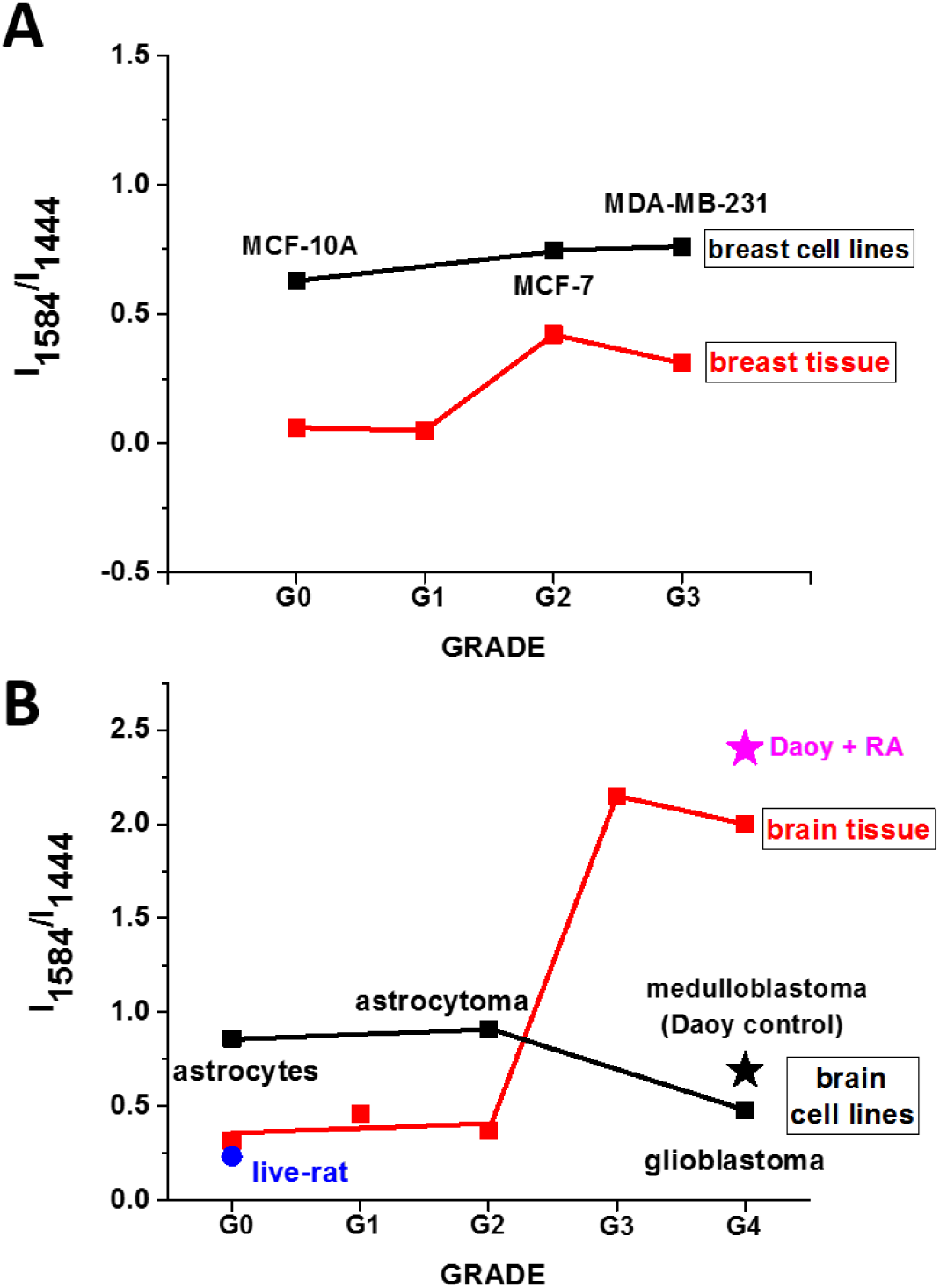
The Raman biomarker of cytochrome c I(1584/1444) as a function of breast cancer grade malignancy G0-G3 ▄ (n=39) (A) and brain tumor grade malignancy ▄ G0-G4 (n=41) (B) at excitation 532 nm, ● in vivo normal rat brain (n=6) (female, Wistar type) at excitation 785 nm, and ▄ in vitro cells at excitation 532 nm of human breast normal cells (MCF-10A) (n=3), malignant (MCF7) (n=3), and agressively malignant (MDA-MD-231) (n=3) (A), and human normal brain astrocytes (NHA) (n=3), astrocytoma (CLR1718) (n=3), glioblastoma (U87 MG) (n=3), and medulloblastoma (Daoy) - control cells ⋆ (n=3), and Daoy+RA ⋆ (n=3) (B).

The sensitivity and specificity for the Raman biomarker I(1584/1444) for tissue breast cancer grade malignancy were obtained directly from Partial least squares discriminant analysis (PLS-DA) and cross-validation and were equal to sensitivity: 83.3% for G0-G1 and 100% for G2-G3 for cross-validation, specificity: 100 % for G0-G1 and 83.3 % for G2-G3 for cross-validation (band at 1584 cm^-1^ corresponds to Cyt c, band at 1444 cm^-1^ corresponds to lipids). Details of statistical analysis are given in the supplementary material. The sensitivity and specificity for the Raman biomarker I(1584/1444) for tissue brain tumor grade malignancy were obtained directly from PLS-DA and cross-validation and were equal to sensitivity: 88.9% for G0-G2 and 100% for G3-G4 for cross-validation, specificity: 100% for G0-G2 and 88,9% for G3-G4 for cross-validation. Details of statistical analysis are given in the Supplementary Material.

In the view of the results presented in Fig.3 it is evident that the Raman biomarker I(1584/1444) based on Cyt c in the human tissues correlates with breast and brain tumour aggressiveness. It indicates that Cyt c plays a crucial role in the development and progression of cancer. The dependence of the Raman biomarker I(1584/1444) of the reduced Cyt c in Fig.3 vs cancer malignancy has a threshold character. The optimal concentration of cytochrome c in the mitochondrium cells that are needed to maintain cellular homeostasis corresponds to the ratio 0.05 for the breast tissues and 0.37 for brain. The concentrations of the reduced Cyt c below the threshold modulate protective, signaling-response pathways, resulting in positive effects on life-history traits. The reduced Cyt c levels above the threshold trigger a toxic runaway process and aggressive cancer development (0.42 for breast and 2.15 for brain). The plot provides an important cell-physiologic response, normally the reduced Cyt c operates at low, basal level in normal cells, but it is strongly induced to very high levels in pathological cancer states by certain cellular stress, the most obvious of which is Reactive Oxygen Species - ROS stress or certain nutrients deficiency.

The second important finding presented in Fig.3 is the striking trend for in vitro cells, which is evidently opposite to that observed in the tissues. The Raman signal I(1584/1444) increases in the breast and brain tissues, whereas it decreases with breast and brain tumour aggressiveness in in vitro cells. This effect is particularly pronounced in the brain (Fig.3B). This finding demonstrates evidence that the tumour environment plays a crucial role in cancer development. During the last decade this conclusion has been solidified and demonstrated that cancer cells must encompass the tumor microenvironment, revealing that the biology of tumors can no longer be understood simply by studying isolated cells in in vitro cultures.^24^ Our results from Fig.3 confirm the importance of tumor microenvironment, which comprises prominent components of cancer progression within the immediate vicinity of tumor cells such as fibroblasts, immune cells and the extracellular matrix.^25,26^ The results demonstrate that there must exist intracellular reductants in tissue other than the reduced nicotinamide adenine dinucleotide (NADH) (present in the growing medium in cell culturing) that contribute to Cyt c activity. Some hints on the essential components from the microenvironment that are important for the activity of Cyt c in the tissues can be provided by detailed analysis of Raman spectra. For this purpose we compared the Raman spectra of the brain tumor tissue using different laser excitation wavelengths. This approach might generate Raman resonance enhancement for some tissue components that cannot be visible for non-resonance conditions.

Raman spectra of lipid droplets in glioblastoma U87MG and in normal astrocytes NHA cells (FigureS2 in supplementary material) in the high frequency region recorded at 355, 532, 785 nm excitations show spectacular differences in Raman spectra due to the various excitation wavelengths. Fig.S2 demonstrates a significant Raman resonance enhancement at 355 nm where the family of retinoids have the absorption.

Until now, vitamin A (retinol) was solely regarded as a biochemical precursor for bioactive retinoids such as retinaldehyde and retinoic acid (RA), but recent results indicate that this compound has its own physiology. It functions as an electron carrier in mitochondria. By electronical coupling between protein kinase Cδ (PCKδ) with Cyt c, vitamin A enables the redox activation of this enzyme.^27^ Fig.S2 demonstrates a significant Raman resonance enhacement at 355nm where the family of retinoids have the absorption (Fig.S3) and methabolism of Vit A.

We wanted to evaluate the possibility and the modality of recovery of cells receiving redox stimuli by retinoic acid (RA) in the absence of cell to cell interactions in in vitro cultures. For these purposes, we incubated the medulloblastoma cells (Daoy) with retinoic acid (RA).

Fig.4 shows microscopy, fluorescence and Raman images of the medulloblastoma (Daoy) cells at the 532 nm wavelength excitation and Raman spectra of nuclei, lipid droplets, mitochondria, cytoplasm and cell membrane obtained from cluster analysis.

**Fig.4.**
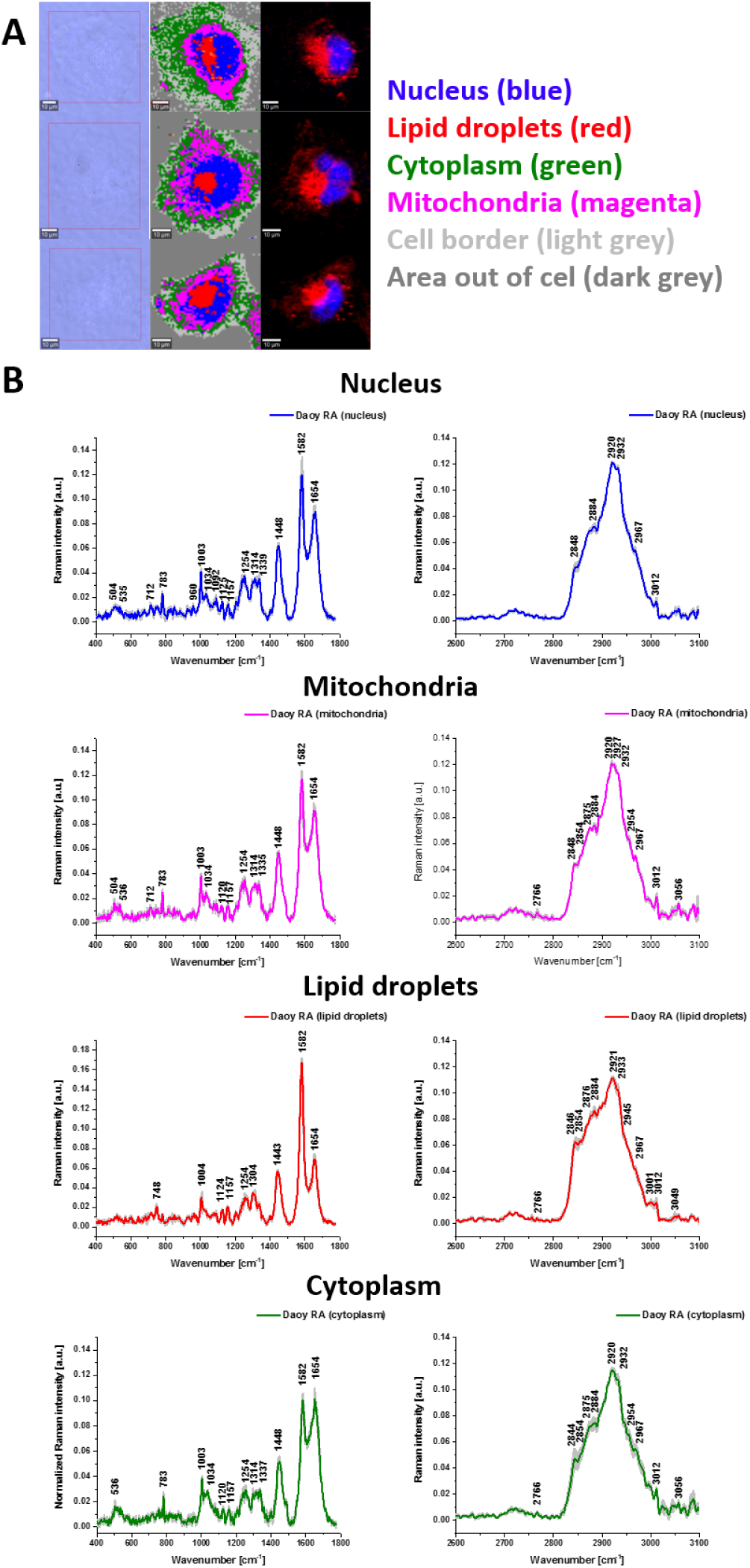
Confocal Raman spectroscopy analysis of the the medulloblastoma (Daoy) cell line incubated with retionoic acid (50μM by 24 hours) at the 532 nm wavelength excitation. (A) Microscopy images; Raman cluster images (nucleus (blue), mitochondria (magenta), lipid droplets (red), cytoplasm (green), cell border (light grey), area out of cell (dark grey)) and fluorescence images of Oil Red O (lipids (red)) and Hoechst 33342 (nulecus (blue)) staining. (B) Mean Raman cluster spectra with SD of nucleus (blue), mitochondria (magenta), lipid droplets (red), cytoplasm (green).

Fig.5 shows the effect of RA on different organelles in the medulloblastoma cells.

**Fig.5.**
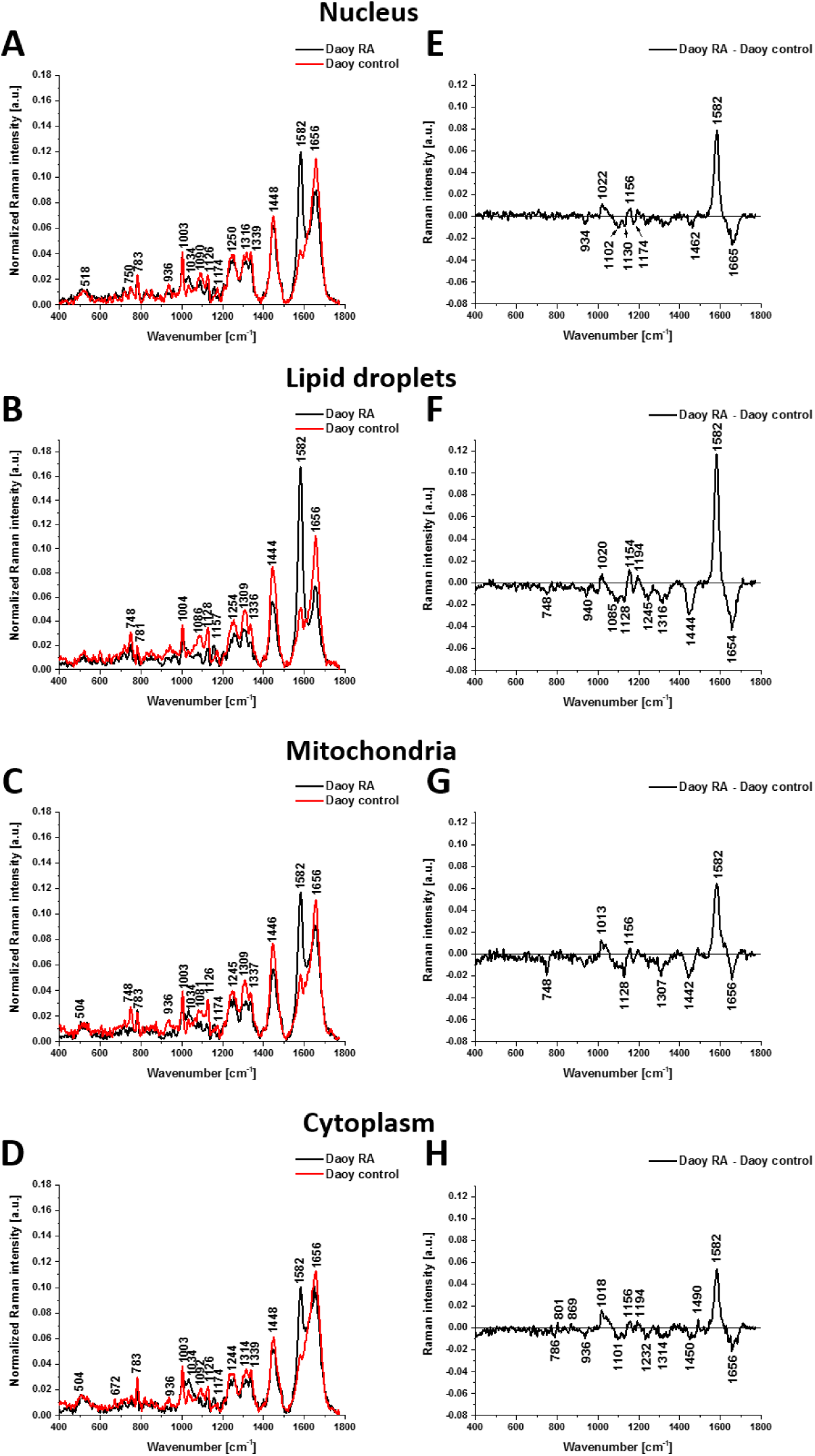
The average Raman spectra obtained at 532 nm wavelength laser excitation from the cluster analysis of nuclei (A), lipid droplets (B), mitochondria (C) and cytoplasm (D) of human medulloblastoma control cells (Daoy) (red line) and Daoy cells incubated with 50 μM of retinoic acid (RA) by 24 hours (black line). Difference spectra of nuclei (E), lipid droplets (F), mitochondria (G) and cytoplasm (H); (n=3).

Interestingly, upon incubation in vitro cells with RA, the reduced Cyt c dramatically increases and reach the level of the Raman signal of the reduced form of Cyt c observed in the brain and breast tissues of aggressive cancers in Fig.3. Indeed, the difference spectra in Fig.5 shows that the intensity of the band at 1584 cm^-1^ of the reduced Cyt c increases upon incubation with RA.

Fig.5 shows the average (n=3) Raman spectra for the nucleous, lipid droplets, mitochondria, and cytoplasm and the alterations upon RA incubation. The results in Fig. 5 show that retinoic acid is an important reducing agent of cytochrome c in human brain cells. Indeed, the intensity of the Raman band at 1584 cm^-1^ increases significantly upon incubation with retinoic acid indicating that the concentration of the reduced Cyt c spectacularly increases. This effect is as strong as in the tumour brain tissue of the high grade G4 (medulloblastoma, glioblastoma) (Fig.3) where the concentration of the reduced Cyt c spectacularly increases.

The results from Fig.4 demonstrate evident coupling between Cyt c and retinoic acid. The mechanism of the coupling between RA and Cyt c in mitochondria is still unknown. The effect is the most visible in mitochondria and lipid droplets (Figs.5B,C). Figs. 5D-F) show that most peaks of Cyts c, b (748, 1128, 1307 and 1337 cm^-1^), lipids (1442 cm^-1^) and α−helix of proteins (1656 cm^-1^) diminished in amplitude following incubation of medulloblastoma cells with RA, whereas the peak at 1584 cm^-1^ increases dramatically upon incubation. The coupling between Cyt c and retinoic acid support previous reports that CYP 26 enzyme, which belongs to the cytochrome family, regulates the cellular level of retinoic acid which is involved in regulation of gene expression.^28^

One can see from Fig.5 that the intensity of the band at 1584 cm^-1^ corresponding to the reduced Cyt c increases drastically upon incubation with RA for all organelles, but the effect is the strongest for lipid droplets. The Raman biomarker I(1584/1444) ratio for mitochondria increases from 0.69 to 2.40 upon incubation with RA and is similar to the values in the brain tissue for G4 (2.00) in Fig.3. These results prove that Raman spectroscopy can indeed probe the reduction of cytochromes in situ.

A major advantage of the Raman imaging is information on distribution of various chemical components in defined cellular organelles in contrast to conventional methods (LC/MS, NMR, and HPLC) that rely on bulk or fractionated analyses of extracted components.

Fig.5 shows the effect of RA on different cell’s organelles. One can see that the difference spectrum at 750 and 781 cm^-1^ (corresponding to cytochrome and DNA vibrations, respectively) is equal to zero for nucleus suggesting that RA does not exert their functions by binding to nuclear receptors. However, the strong Raman signal at 1584 cm^-1^ of reduced Cyt c indicates that the genomic action with induction or repression of the expression of target genes plays a role. This finding seems to support the suggestions that RA bind to cellular retinoic acid binding protein (CRABP) in cytosol and that this complex migrates to nucleus to exert its effects through binding to nuclear receptors to retinoic acid (RAR or RXR).^29,30^

The results in Fig.5 shows the significant effect of cytochrome activity on lipid droplets (Fig.5B) and mitochondria (Fig.5C) where Cyt c is widely believed to be localized solely in the mitochondrial intermembrane space under normal physiological conditions.^31^

The significant effect of cytochrome activity on lipid droplets (Fig.5B) and mitochondria (Fig.5C) suggests that nongenomic action though regulation of signaling pathways dominates. Fig.2 provides hints related to the coupling between the haem functional center and the tyrosine residual in Cyt c. Indeed, at 785 nm excitation when the Cyt c resonance enhancement disappears the vibrational modes of tyrosine and phosphorylated tyrosine are revealed.^22^ The Raman resonance enhancement at 785 nm may be due to absorption zinc-finger-like activation domain at around 700 nm.^27^

To elucidate whether Cyt c is released from mitochondria to diffuse in the cytosol we studied the concentration of Cyt c in cytosol. Note, that our results in Fig.5D shows the Cyt c activity in the cytoplasm, which indicates that we can measure the amount of Cyt c leaking from mitochondria to cytosol. Indeed, the Raman peak at 1584 cm^-1^ for NHA which is slightly blue shifted to 1582 cm^-1^ for cancerous cell lines (CRL1718, U87MG and Daoy) in Fig.5D demonstrates the presence of Cyt c in cytosol. The release of Cyt c from mitochondria to the cytosol activates apoptosis by the stream of caspase and proteases.^32–34^ The concept that programmed cell death by apoptosis serves as a natural barrier to cancer development.^24^ Therefore, the results provide new tools to check the role of Cyt c in an anti-apoptotic switch by phosphorylation of Tyr48.^35^

One can see that the Raman signal at 1307 cm^-1^ increases in agressive tumor (Daoy) indicating that the Cyt c has been released to cytoplasm.

The second important point is that the Raman intensity of the 1337 cm^-1^ vibrational peak corresponding to cytochrome b-type does not change in the difference spectrum (Fig.5D-F) upon incubation with RA, which indicates that electrons from RA are not transfered to the complex III, but directly to Cyt c. The mechanism is unknown, but it was suggested that retinol functions as an electron carrier in mitochondria by electronically coupling protein kinase Cδ (PCKδ) with Cyt c, and vitamin A enables the redox activation of this enzyme. In silico docking studies confirmed other vitamin A metabolites have similar binding capacity as retinol itself: they possess the same β-ionone structure, while being chemically distinguished by modifications of their polyene tails.^27^

## Discussion

Fig.6 shows that concentration of the reduced cytochrome c (estimated from the intensity of the 1584 cm^-1^ band) in the living cells under in vitro conditions where the interaction with the environment has been eliminated decreases with brain tumor aggressiveness. The Cyt c is localized in the inner membrane of mitochondrium and transfers electrons between complex III and complex IV of the respiratory chain (Fig.6E) generating the proton gradient and triggering ATP production. The heme chromophore of Cyt c accepts electrons from the bc1 complex (complex III) and transfers electrons to the complex IV. Under normal physiological conditions, when the electron transport chain and chemiosmosis make up oxidative phosphorylation, there is a balance between the oxidized and reduced forms of Cyt c, which depends on the efficiency of the electron transport from complex III which produces the reduced form of Cyt c and the transfer of electrons from Cyt c to complex IV resulting in producing the oxidized form of Cyt c. Under cancer development a balance between the oxidized and reduced forms of Cyt c is broken (or reduced) and we observe a lower pool of Cyt c in the reduced form (and a larger pool of cyt c). The lower concentration of the reduced form of Cyt c in cancer cells in vitro when compared with the normal cells as presented in Fig.6 indicates that the balance between the reduced and oxidized forms is deregulated. To learn how this balance is deregulated we analyzed the Raman biomarker of the Cyt b at 1337 cm^-1^ that dominates the complex III (Fig.6E) that transfers electrons to Cyt c. Fig.6 shows that the amount of the Cyt b decreases vs tumor malignancy, which indicates sudden limitation of the electron flow from the complex III to Cyt c resulting in less effective production of the reduced form of Cyt c (see Fig.6A,B at 1584 cm^-1^). It indicates that the reaction catalyzed by ubiquinol-cytochrome-c reductase between quinol (QH_2_) and ferri- (Fe^3+^) Cyt c, QH_2_ + 2 ferricytochrome c Q + 2 ferrocytochrome c + 2 H^+^ is less effective to produce quinone (Q), ferro - (Fe^2+^) Cyt c, and H^+^ and the tumor cells lose the respiratory function. The mechanism of deficiency in the complex III subunit Cyt b is unknown.

**Fig.6.**
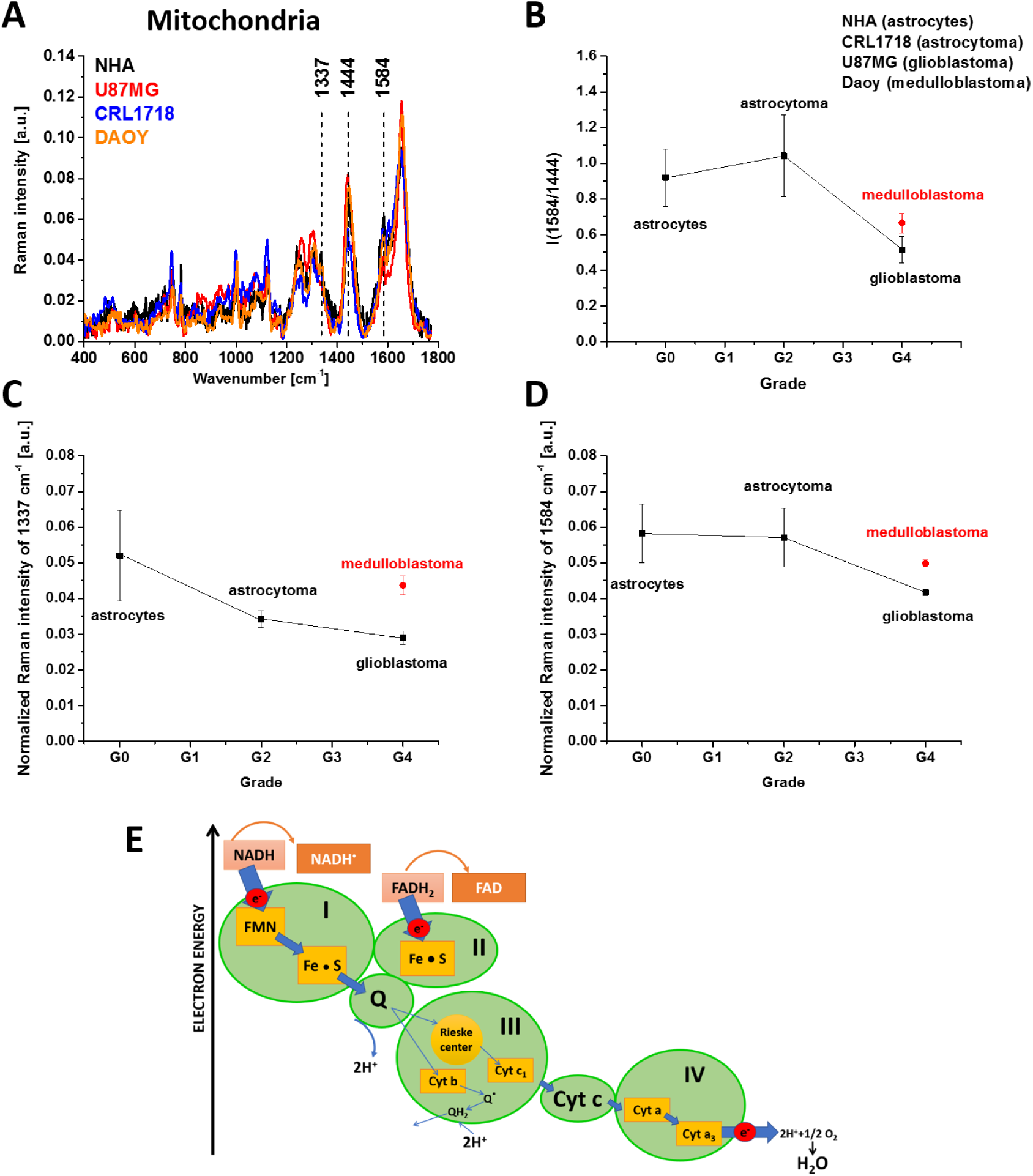
Normalized mean cluster Raman spectra of mitochondria of human brain astrocytes (NHA), astrocytoma (CRL1718), glioblastoma (U87MG) and medulloblastoma (Daoy) cells (A) and the concentration of the reduced cytochrome c (estimated from the intensity of the 1584 cm^-1^ band) in the cells under in vitro conditions where the interaction with the environment has been eliminated in a function of brain tumor aggressiveness (B). (C) The amount of the cytochrome b based on the normalized Raman intensity of the peak 1337 cm^-1^ and (D) reduced cytochrome c (peak at 1854 cm^-1^) vs human brain tumor malignancy. (E) The respiratory chain generating the proton gradient and triggering ATP production.

Our results demonstrate that the mechanisms controlling the electron transport chain are spectacularly deregulated in cancers and indicate that electron transport, organized in terms of electronegativity, is inhibited between complex III and Cyt c (Fig.6E). It may be associated with mitochondrial complex III deficiency in electron chain transfer in cancers. Our results support earlier suggestions that the Qo site of the mitochondrial complex III is required for the transduction of hypoxic signaling via reactive oxygen species production.^36^

Cancer cells deficient in the complex III subunit Cyt b, which are unable to maintain respiratory function, increase ROS levels and stabilize the HIF-1α protein during hypoxia.^36^

The deficiency of mitochondrial complex III in electron chain transfer in cancers may be related to conformational changes clearly visible in the region of the heme ruffling deformation directly tied to the protein induced distortion^37–39^ and 1126 cm^-1^ (vibrations of C_b_-CH_3_ side radicals) is reflected by shift in the position of the peak maxima.

## Conclusions

This paper presents a non-invasive approach to study redox status of cytochromes *in vitro* human brain cells of normal astrocytes (NHA), astrocytoma (CRL-1718), glioblastoma (U87- MG) and medulloblastoma (Daoy), and human breast cells of normal cells (MCF10A), slightly malignant cells (MCF7) and highly aggressive cells (MDA-MB-231), in vivo animal models, and *ex vivo* brain and breast tissues surgically resected human specimens by means of Raman microspectroscopy at 355 nm, 532 nm, 785 nm and endospectroscopic Raman probe at 785 nm.

We showed that the amount of reduced cytochrome c (at 1584 cm^-1^) is lower in cancer cells when comparted with the normal cells at in vitro conditions when the effect of microenvironment is eliminated. Incubation in vitro with RA increases the amount of reduced Cyt c to the level observed in cancer cells in tissues. In the tissue, where retinoids are delivered to the organism with diet, and the microenvironment interacts with the cancer cells the amount of reduced Cyt c (at 1584 cm^-1^) becomes abnormally high, much higher than in normal cells, particularly for the brain tumour (for G0 = 0.306, for G4 = 1.992)

Our results demonstrate that the mechanisms controlling the electron transport chain are spectacularly deregulated in cancers and indicate that electron transport, organized in terms of electronegativity, is inhibited between complex III and Cyt c. It might be associated with mitochondrial complex III deficiency in electron chain transfer in cancers.

This study sheds light on a largely uninvestigated triangle between cytochromes, lipid metabolism and mitochondrial function. Insight into this organelle crosstalk increases our understanding of mitochondria-driven cell death. Our findings furthermore provide a first hint on the role of Cyt c in mechanisms that regulate cancer progression by using redox-sensitive mitochondrial cytochrome Raman bands for label-free detection of mitochondrial dysfunction.

## Methods

### Ethics statement

All the conducted studies were approved by the local Bioethical Committee at the Polish Mother’s Memorial Hospital Research Institute in Lodz (53/216) and by the institutional Bioethical Committee at the Medical University of Lodz, Poland (RNN/323/17/KE/17/10/2017). Written consents from patients or from legal guardians of patients were obtained. All the experiments were carried out in accordance with Good Clinical Practice and with the ethical principles of the Declaration of Helsinki. Spectroscopic analysis did not affect the scope of surgery and course and type of undertaken hospital treatment.

### Patients

In the presented studies the total number of patients diagnosed with brain tumors was 41. Among them 11 were diagnosed with medulloblastoma, 1 with embryonic tumor PNS, 3 with ependynoma anaplastic, 4 with ependymoma, 2 with astrcytoma fibrous, 1 with astrocytoma, 1 with ganglioma, 8 with astrocytoma pilocytic, 1 with subependymoma, 2 with hemangioblastoma, 4 with craniopharyngioma, 1 with dysembryoplastic neuroepithelial tumor, 1 with papillary glioneuronal tumor and 1 sample it was tumor metastasis. All patients were treated at the Polish Mother’s Memorial Hospital Research Institute in Lodz. For breast cancers the number of patients was 39, all patients were diagnosed with infiltrating ductal carcinoma and treated at the M. Copernicus Voivodeship Multi-Specialist Center for Oncology and Traumatology in Lodz.

### Tissues samples collection and preparation for Raman spectroscopy

Tissue samples were collected during routine surgery. The non-fixed samples were used to prepare 16 micrometers sections placed on CaF_2_ substrate for Raman analysis. In parallel typical histopathological analysis by professional pathologists from the Polish Mother’s Memorial Hospital Research Institute in Lodz for brain tissues samples or from Medical University of Lodz, Department of Pathology, Chair of Oncology for breast tissues samples was performed. The types and grades of tumors according to the criteria of the Current WHO Classification were diagnosed.

### Laboratory animals breeding and in-vivo experiments protocols

Laboratory animals used in *in-vivo* experiments were bred in a certified unit - the laboratory of the Faculty of Biology and Environmental Protection of the University of Lodz in Lodz, identification number 10616205 (certificate of the poviat veterinary inspector No. 7/4617/2007 of 05 July 2007 about meeting the required conditions breeding and maintenance of experimental animals, decision of the poviat veterinary officer No. 248/z/2007 of August 9, 2007 on assigning a veterinary identification number, decision of the Minister of Education and Science No. 18/2005 of December 21, 2005 on entering the unit to the list of entities authorized to conduct animal experiments). Six female WISTAR [CV-I] rats were prepared for the experiment, F11/2 generation, white light, outbred herd, own breeding, mass 200-230 g. Before starting the experiment, animals were provided with adaptation and acclimatization for a period of 4 days, while maintaining standardization of environmental factors in the breeding and later during the experiment. Before the in-vivo experiments, the veterinarian assessed the animal’s state of health and condition after the acclimatization period, and before the procedure and introduction of the animal under general anesthesia. Particular attention was paid to: animal appearance (smooth, dry coat, without dirt and secretions around the eyes, nose and anus, skin in unpigmented places pink), animal behavior (active animal, shows interest in the environment), appetite, water intake (does not show a lack of appetite, increased water intake or a lack of water intake), animal body weight (normal), breathing method (no deviations), hydration level, assessed by the speed of the skin’s recovery to its previous state (no dehydration of the body).

Anesthesia protocol: before performing the procedure, the following was prepared: the order of operations was planned and the necessary equipment prepared - including surgical instruments, needles, syringes; equipment needed for the experiment was checked; drugs used for euthanasia and sacrifice were prepared (the date of evaluation was checked, appropriate drug uses were selected). Injected anesthesia into the intraperitoneal space (i.p.) with a mixture of ketamine + xylazine at a dose of the active substance of 5/10 + 40/80 mg / kg was used. After anesthesia, the depth of anesthesia was monitored: eyelid reflex checked – atrophy; and corneal - partially preserved; breathing - regular breaths, evenly deep, frequency slightly lower than in the state of consciousness; coloration of mucous membranes - pink, moist. During anesthesia, the correct body temperature of the animal was also maintained, protecting it from hypothermia by heating under a heating lamp and covering sensitive areas of the body. 0.9% saline was administered into the conjunctival sac.

Animals were sacrificed for organ removal by an overdose of the anesthetic agent - sodium pentobarbital using a detailed assessment of euthanasia methods based on the monograph “Recommendations for euthanasia ofexperimental animals” Laboratory Animals Ltd. The organs were collected after the confirmation of death features: cardiac and respiratory arrest, no reflexes, reducing the body temperature below 25°C. The animals bodies after experiments were secured and sent for utilization in accordance with the internal procedure: Agreement No. 2/2019 the company EKO-ABC with headquarters in Bełchatów, Poland.

### Cell culture and preparation for Raman spectroscopy

The studies were performed on a normal human astrocytes (Clonetics NHA), human astrocytoma CCF-STTG1 (ATTC CRL-1718) and human glioblastoma cell line U87MG (ATCC HTB-14) purchased from Lonza (Lonza Walkersville. Inc.) and American Type Culture Collection (ATCC), respectively. The NHA cells were maintained in Astrocyte Medium Bulletkit Clonetics (AGM BulletKit, Lonza CC-3186) and ReagentPack (Lonza CC-5034) without antibiotics in a humidified incubator at 37°C and 5% CO_2_ atmosphere. The U87MG cells were maintained in Eagle’s Minimal Essential Medium Eagle with L-glutamine (ATCC 30-2003) supplemented with 10% fetal bovine serum (ATCC 30-2020) without antibiotics in a humidified incubator at 37°C and 5% CO_2_ atmosphere. The CRL-1718 cells were maintained in RPMI1640 Medium (ATCC 302001) supplemented with 10% fetal bovine serum (ATCC 30- 2020) without antibiotics in a humidified incubator at 37°C and 5% CO_2_ atmosphere.A human desmoplastic cerebellar medulloblastoma cell line (ATCC HTB-186, Daoy) was grown in Eagle’s Minimum Essential Medium (EMEM, ATCC 30-2003) supplemented with the fetal bovine serum to a final concentration of 10% (Gibco, Life Technologies, 16000-044). Cells were maintained without antibiotics at 37°C in a humidified atmosphere containing 5% CO_2_.

Cells were seeded on CaF_2_ window in 35mm Petri dish at a density of 5×10^4^ cells per Petri dish the day before examination. Before Raman examination, cells were fixed with 4% formalin solution (neutrally buffered) and kept in phosphate buffered saline (PBS, Gibco no. 10010023) during the experiment. After the Raman imaging measurements, the cells were exposed to Hoechst 33342 (25 μL at 1 μg/mL per mL of PBS) and Oil Red O (10 μL of 0.5 mM Oil Red dissolved in 60% isopropanol/dH_2_O per each mL of PBS) by incubation for 15 min. The cells were then washed with PBS, followed by the addition of fresh PBS for fluorescence imaging on an Alpha 300RSA WITec microscope.

For the experiment with retinoic acid supplementation, the Daoy cells were incubated with 50 μM of RA for 24 hours.

### Raman human tissues spectroscopic measurements *ex-vivo*

WITec (Ulm, Germany) alpha 300 RSA+ confocal microscope was used to record Raman spectra and imaging. The configuration of experimental set up was as follows: the diameter of fiber: 50 μm for 355 and 532 nm and 100 μm for 785 nm, a monochromator Acton-SP-2300i and a CCD camera Andor Newton DU970-UVB-353 for 355 and 532 nm, and an Ultra High Throughput Spectrometer (UHTS 300) and CCD camera Andor Newton iDU401A-BR-DD- 352 for 785 nm.. Excitation lines were focused on the sample through a 40x dry objective (Nikon, objective type CFI Plan Fluor C ELWD DIC-M, numerical aperture (NA) of 0.60 and a 3.6–2.8 mm working distance). The average laser excitation power was 1 mW for 355 nm, 10 mW for 532 and 80 mW for 785nm, with an integration time of 0.5 sec, 0.5 sec and 0.5 sec respectively. An edge filters were used to remove the Rayleigh scattered light. A piezoelectric table was used to record Raman images. No pre-treatment of the samples was necessary before Raman measurements. The cosmic rays were removed from each Raman spectrum (model: filter size: 2, dynamic factor: 10) and the smoothing procedure: Savitzky–Golay method was also implemented (model: order: 4, derivative: 0). Data acquisition and processing were performed using WITec Project Plus software.

### Raman rat tissues spectroscopic measurements in-vivo

EmVision HT Raman spectrometer with volume phase holographic transmission grating, combine with an Andor iVac CCD camera and equipped with high efficiency fiber optic probe was used for in-vivo measurements. The excitation wavelength was 785 nm for low frequency region. The probe used during measurements had 2.1 mm outer diameter containing 7 low- hydroxyl, 300 µm core fibers for scattered light collection located around a central ca.300 µm core source fiber. The signal emitted from the sample passed through a ring-shaped notch filter to remove elastically backscattered light and light scattered from the lens surface for 785 nm excitation laser line. A sapphire lens was used at the tip of the probe to overlap the excitation and detection volumes. The optic probe collection fibers was connected to the spectrometer through a single SMA connector. The laser spot size determining tissue excitation area had a diameter ca. 0.5 mm. The data processing: smoothing, background subtraction, Raman peaks analysis was performed using Origin software.

### Statistical Analysis

Statistical significance was verified by analysis of variance, employing the Fisher’s Least Significant Difference test. The mean and standard deviation (SD) were calculated for all of the investigated parameters. The data were analyzed using Origin software for Windows. Significant differences between the treatment groups and controls were compared using one-way analysis of variance (ANOVA); Data with p-values that were lower than 0.05 were identified as statistically significant.

The data were processed making use of multivariate analysis. Data processing was performed using Project Plus (WITec GmbH, Germany), Origin 2016 (OriginLab, USA) and MATLAB (MathWorks, USA) with PLS-Toolbox (EigenvectorResearch Inc., USA).

## Data availability

The request for data sets, both raw and processed data, generated during the present study can be agreed and made directly to the corresponding author.

## Author Contributions

Conceptualization: HA, JS, BB-P; Funding acquisition: HA, JS; Investigation: BB-P, JS; Methodology: HA, BB-P, JS; Writing – original draft: HA; Writing – review & editing: HA, JS, BB-P. All authors reviewed and provide feedback on the manuscripts.

