## Supplementary material for "Redox state changes of mitochondrial cytochromes in brain and breast cancers by Raman spectroscopy and imaging": Suplementary Material

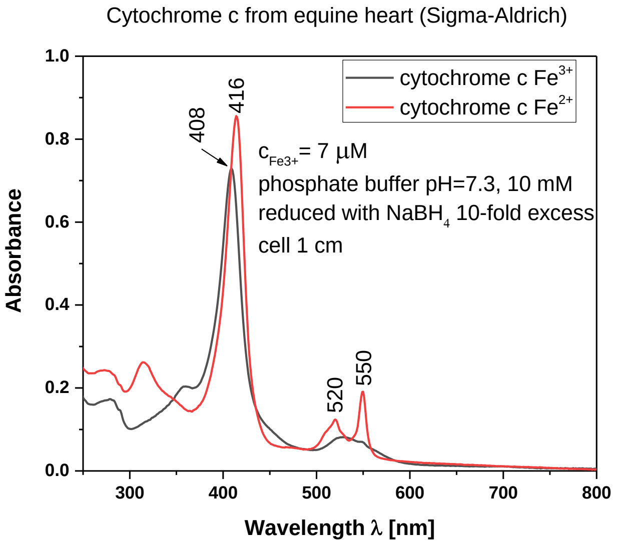


**Figure S1.** Electronic absorption spectra of cytochrome c in ferric and ferrous states. Ferrous cytochrome c was prepared by adding 10-fold excess $NaBH_4$ (as a reductor).

Figure S2 shows Raman spectra of lipid droplets in glioblastoma U87MG cells (Fig.S2A) and in normal astrocytes NHA cells (Fig.S2B) in the high frequency region characteristic for the $CH_2$ and $CH_3$ stretching vibrations of lipids, DNA and proteins at 355, 532, 785 nm excitations. One can see spectacular differences in Raman spectra due to the various excitation wavelengths. Fig.S2 demonstrates a significant Raman resonance enhancement at 355 nm where the family of retinoids have the absorption (Fig.S3).

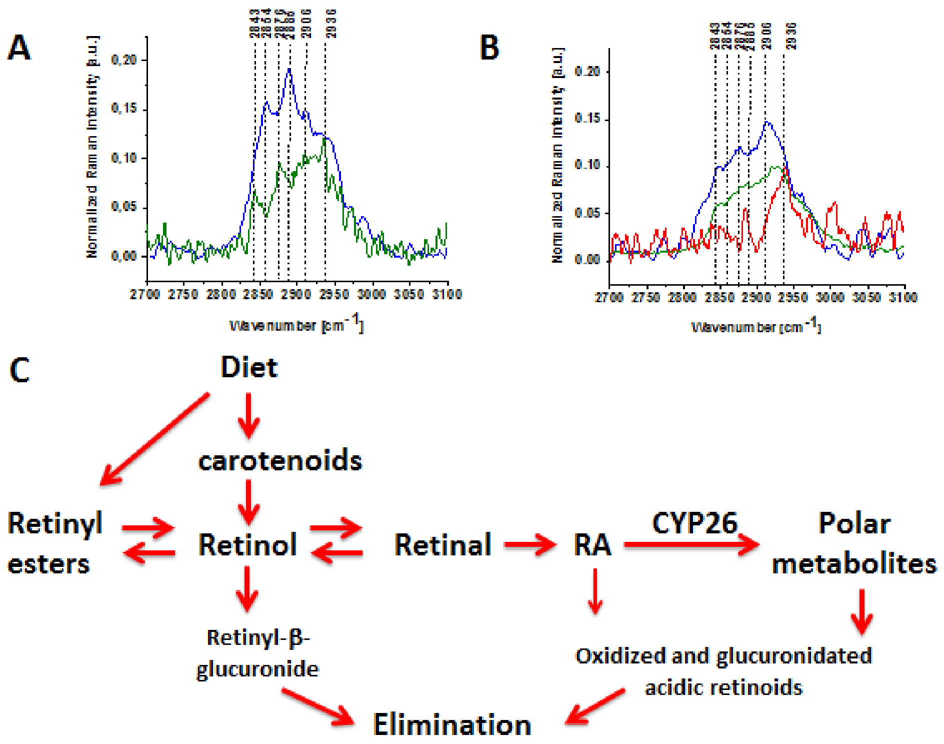


**Figure S2.** Raman spectra of lipid droplets in glioblastoma U87MG (A) and in normal astrocytes NHA cells (B), ▬ 355 nm (blue), ▬532 nm (green), ▬780 nm (red), metabolism of vitamin A (C).

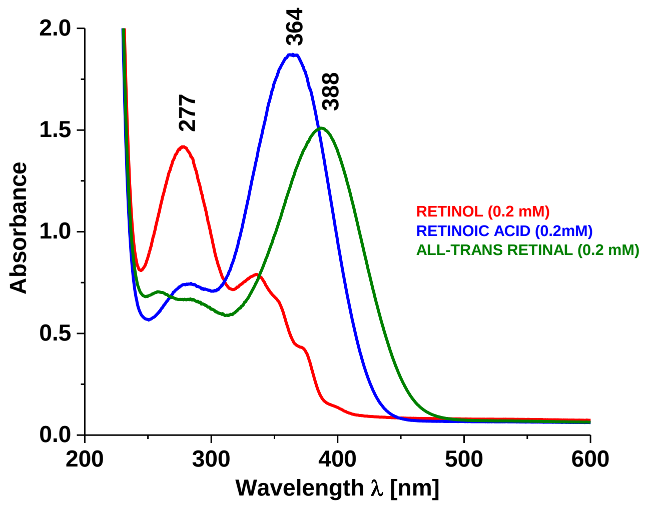


**Figure S3.** Absorption spectra of retinoids. Retinol (c=0.2 mM in chloroform, red), retinoic acid (c=0.2 mM in chloroform, blue), all-trans retinal (c=0.2 mM in chloroform, green); with a cuvette path length of 1 mm.

**Statistical Analysis - continued**

We have used a hierarchical cluster analysis (HCA) of confocal Raman dataset of individual cells to construct Raman spectra images of chemically different regions. Briefly, HCA is an unsupervised statistical method that clusters data according to their similarities. The HCA helps to determine different features in a Raman dataset where spectral distances are calculated and each Raman spectra are assigned to clusters based on these distances.

The sensitivity and specificity for calibration and cross-validation data sets were calculated using PLS-DA (Partial least squares discriminant analysis) method implemented in MATLAB. PLS-DA was used to find and illustrate the maximum covariance between input data and predefined information about belonging to a given class. Similarly as in PCA (Principal Component analysis) it was done by linear transformation of the input data to a new system of orthogonal coordinates so-called latent components. Traditionally, the first component reflected maximum variation between classes. Briefly, during PLS-DA analysis input data are in a form of x matrix with dimensions m x n, where each of the m lines represents a single sample (spectrum) described by n variables (wavenumbers). Information on belonging to a given class is coded in the zero-one matrix Y (m x k-1), where k is the number of classes. The X data in the new coordinate system is expressed by their linear combination with the W* (X-weight) weight matrix. whose columns {w * i, ..., w * m} are a set of new orthogonal base vectors $T=XW^*$.As in PC A, case factor coordinates multiplied by factor loadings are a good approximation of X: $X=TP^T+E$, in addition, multiplied by the weights C (Y-weight) predict the value of Y according to the formula: $Y=TC^T+F$ where E and F are residual matrices. Finally, Y = XW*CT + F = XB + F, where matrix B is a matrix of PLS regression coefficients reflecting the relationship between X and Y. If we suppose that $C_x$ = cov(X ) = (1/n-1) $X^TX$, $C_y$ = cov(Y ) = (1/n-1) $Y^TY$ and $C_{XY}$ = cov(XY) = (1/n-1) $X^TY$ will be the X and Y covariance matrices, and the covariance between X and Y, respectively for previously centered X and Y matrices, than to determine vectors w* in PLS methods we can use the NIPALS algorithm (nonlinear iterative partial least squares) based on the presentation of the association of covariance with the Pearson correlation coefficient sing the formula:$[C_{XY}]^2$ =[cov(X Y]$^2$ = cov(Xa)[corr(XaYb)]$^2$ cov(Yb). Thus PLS can be treated as a form of canonical correlation analysis (CCA), in which the criterion of maximum correlation is balanced by the simultaneous request to find the highest possible variance in both X and Y data. Because in PLS-DA, the variance of data in the class matrix (Y) is not significant to solve the problem we have to solve the equation: (1/n-1) $X^T$ $Y(YY^T)^{-1}Y^TXw^*=Hw^*$, whose solution is a vector $w^*$ corresponding to the highest self-value matrix H composed of sums of squares between classes. It should be noted that PLS-DA is an iterative method, that's why to calculate the next vector w* the residual matrix is created, which is reduced by one order of the matrix X, and the calculations are repeated so long as the X order of the matrix is exhausted.

In our PLS-DA model sensitivity is a true positive rate that measures the proportion of actual positives that are correctly identified, while specificity, called the true negative rate measures the proportion of actual negatives that are correctly identified. In our case the correctly identified samples there were samples for which the ratio of Raman peaks intensities was adequate to the ratio typical for a given grade of brain tumor or breast cancer.

The sensitivity and specificity for brain tumor were obtained directly from PLS-DA and cross-validation and were equal to:

sensitivity: 100% for G0-G2 and 100% for G3-G4 for calibration

specificity: 100% for G0-G2 and 100% for G3-G4 for calibration

sensitivity average value: 100%, specificity average value: 100% for calibration

sensitivity: 88.9% for G0-G2 and 100% for G3-G4 for cross-validation

specificity: 100% for G0-G2 and 88,9% for G3-G4 for cross-validation

sensitivity average value: 94.45%, specificity average value: 94.45% for cross-validation

The sensitivity and specificity for breast cancer were obtained directly from PLS-DA and cross validation and were equal to:

sensitivity: 100% for G0-G1 and 100% for G2-G3 for calibration

specificity: 100% for G0-G1 and 100% for G2-G3 for calibration

sensitivity average value: 100%, specificity average value: 100% for calibration

sensitivity: 83,3% for G0-G1 and 100% for G2-G3 for cross validation

specificity: 100% for G0-G1 and 83,3% for G2-G3 for cross validation

sensitivity average value: 91.7%, specificity average value: 91.7% for cross vali-dation

For Raman biomarker I(1586/1444) (for a ratio of band intensities 1586/1444) in spectra with a mixed fat-protein profile from the entire imaging area of human brain tumor and breast cancer tissue at 532 nm excitation and normal rat brain tissue (type Wistar) at 785 nm excitation, the Kruskal-Wallis test at the confidence level $p \leq 0.05$ has confirmed that the values of this biomarker for particular grades of brain tumors belong to statistically different groups.

The established Raman biomarker values for individual brain tumor and breast cancer malignancy grades are shown in Table 1 with SD - standard deviation.

**Table 1**

| Malignancy tumor grade | Brain tumor | | Breast cancer | |
|---|---|---|---|---|
| | Average value | SD | Average value | SD |
| G4 | 2.00 | 1.03 | - | - |
| G3 | 2.15 | 0.99 | 0.31 | 0.16 |
| G2 | 0.37 | 0.09 | 0.42 | 0.27 |
| G1 | 0.46 | 0.16 | 0.05 | 0.004 |
| G0 | 0.32 | 0.18 | 0.06 | 0.03 |
